# A Conditional Degron Tag for RNA Sensing in Bacteria

**DOI:** 10.1101/2024.12.16.628724

**Authors:** Tyler J. Simons, Ming C. Hammond

## Abstract

Here we report the development of a conditional degron system for use in bacteria in which a tagged fluorescent protein (FP) reporter, mVenus-S1, is stabilized only in the presence of TAR RNA. The functional design takes advantage of the bipartite nature of the ssrA degron sequence, so that TAR RNA binding to the inserted Tat peptide sequence blocks degron recognition to rescue protein activity. Several fluorescent proteins and a chemiluminescent enzyme can be tagged and activated using this system, and our results reveal that chromophore maturation time correlates with fold change, with up to ∼60-fold turn-on. Finally, we designed a functional biosensor for *oxyS* sRNA and integrated it into a single transcript in which the FP and RNA-based biosensor are encoded on the same mRNA. To our knowledge, this is the first proof-of-concept for a novel biosensor design that expresses both protein and RNA components from a single transcript. The streamlined system is fully genetically encoded, does not require an exogenous fluorophore, and permits both components to be coordinated by expression from the same promoter.

## INTRODUCTION

The demonstration that small molecule drugs such as PROTACs and molecular glues can induce targeted protein degradation for therapeutic outcomes^1,2^ has inspired the development of degradation tags as tools for mammalian cell biology. Using either natural or engineered degrons, which are specific amino acid sequences that lead to protein degradation typically through the ubiquitin-proteasome system, many small molecule-triggered protein inactivation (AID, HaloPROTAC, dTAG) and protein activation (DD) systems have been described.^3^ . For light-triggered protein inactivation, photoactivatable PROTACs have been developed.^4,5^ For RNA and DNA imaging, a conditional degron system was developed by overlapping the TAR RNA-binding Tat peptide sequence with the C-terminal degron sequence, such that a fluorescent protein tagged with this Tat-degron peptide would be stabilized in the presence of RNAs tagged with the TAR sequence.^6^ This system has been applied in mammalian cells to detect *survivin* mRNA and lncRNA MALAT1^7^ and to visualize genome dynamics through engineering of the sgRNA to make fluorogenic CRISPR.^8^

Recently, conditional degron systems also have been explored for use in bacteria. Deletion of the native SspB chaperone protein in *E. coli*, which binds and delivers substrates to the ClpXP proteasome, generates a strain that degrades degron-tagged proteins controlled by an inducible copy of SspB.^9^ For optogenetic control, a light-activatable split TEV protease called TEVp was applied to process two types of appended degron tags for improving shikimic acid production in *E. coli*.^10^ For light-triggered protein inactivation, TEVp cleaves an N-terminal sequence to reveal a cryptic N-terminal degron. For light-triggered protein activation, TEVp cleaves off a C-terminal degron. A different design for light-triggered protein inactivation is the LOVdeg system, which uses the light-dependent unfolding of the *As*LOV2 domain to control access of the C-terminal degron.^11^ An 11.5-fold decrease in mCherry-*As*LOV2 fluorescence was observed after blue light treatment. In addition, diverse protein targets including the LacI repressor, CRISPRa activator, AcrB efflux pump, and a fatty acid metabolism enzyme were able to be inactivated with blue light.

In this study, we present the development of a conditional degron system for RNA sensing in bacteria (**Fig. 1A**). A novel split degron approach permitted us to adapt the TAR-Tat RNA-peptide interaction for RNA-triggered protein activation in *E. coli*. A variety of fluorescent and chemiluminescent proteins were shown to be robustly activated in the presence of a trigger RNA, and our experiments revealed a strong effect for chromophore maturation rate. In addition, we showed that the protein-RNA pair can be encoded as a single transcript for compactness and designed a TAR-based biosensor to detect a regulatory small RNA (sRNA), *oxyS*.

**Figure 1.**
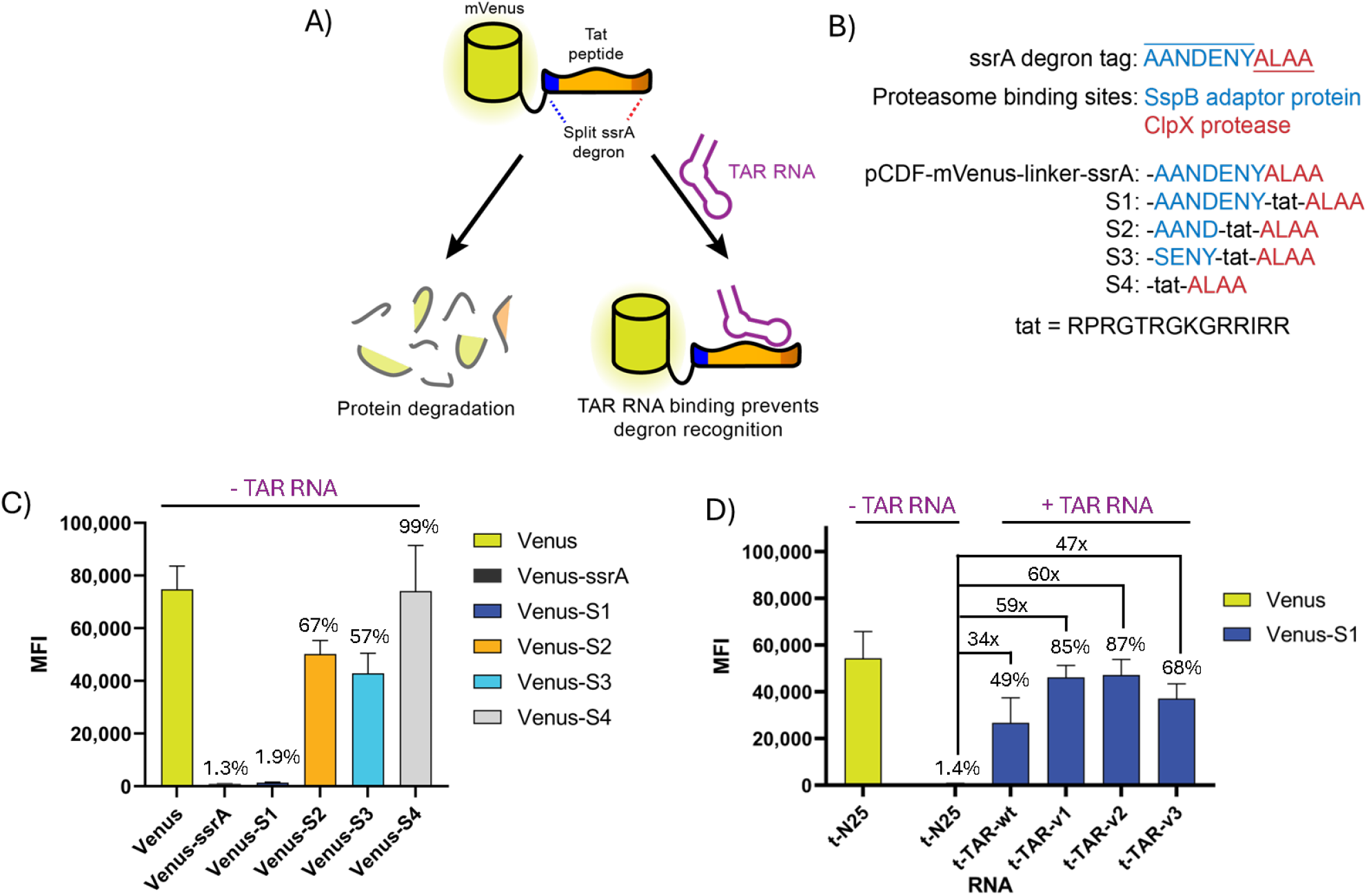
Design and screening of conditional degron constructs for protein degradation and rescue in live bacteria. (A) A conditional degron system using a C-terminal degron tag and TAR RNA. (B) Design and sequences of C-terminal degron tags. (C) Flow cytometry data for *E. coli* expressing mVenus constructs with different C-terminal degron tags. (D) Flow cytometry data for *E. coli* co-expressing mVenus-S1 and different TAR RNA constructs or N25 control in a tRNA scaffold (t). All mean fluorescence intensity (MFI) values were analyzed for at least 30,000 cells per sample and standard deviation is shown for at least three biological replicates. Percentages are relative to mVenus. RNA sequences used in this work are shown in **Supplementary Table 1**.

## RESULTS

The mammalian conditional degron tag was designed by overlapping the arginine-rich BIV Tat peptide, which binds TAR RNA, with the C-terminal degron sequence, RRRG.^6^ This destabilizing mammalian degron was engineered *de novo*^*12*^ and leads to ubiquitination of the tagged protein, causing proteasomal degradation; however, the ubiquitin-proteasome pathway is absent in bacteria requiring a different degron sequence to be implemented.^13^ Initially, a bacterial degron sequence from *E. coli* Mu transposase (MuA), RRKKAI,^14^ was analyzed because it should permit a similar Arg-rich overlap with Tat for *E. coli*. However, mVenus fluorescence was not diminished when the MuA sequence was fused to the C-terminus (data not shown), which supports prior work that other sequence determinants are required for MuA degradation.^15^ Thus, the bacterial conditional degron tag required a different design approach.

The ssrA C-terminal degron sequence, AADENYALAA, is part of the highly conserved transfer-messenger RNA (tmRNA) degradation pathway used to rescue ribosomes stalled on mRNAs in bacteria. Through the activity of tmRNA (also referred to as 10Sa RNA)^16^ to initiate *trans*-translation, the ssrA degron sequence is typically appended to protein products that are missing the stop codon. Besides rescuing ribosomes that were stalled on the stop-less mRNA, the ssrA-tagged protein is degraded by ClpXP, Lon+, and FtsH.^17–20^

Recent literature has revealed two binding motifs within the ssrA degron tag: the first seven amino acids, AANDENY, are bound by the chaperone protein SspB, which binds and delivers substrates to ClpXP, while the final four amino acids, ALAA, are bound by ClpX unfoldase.^17,18^ One study demonstrated that only the final ALAA sequence is required to degrade tagged GFP significantly.^21^ Inspired by these prior findings, we designed three peptide tags, S1-S3, in which the Tat peptide was inserted between the two binding motifs, with the concept that TAR binding could conceal one or both motifs to block degradation (**Fig. 1B**). The S4 tag has Tat peptide before the minimal degron tag without the SspB binding motif. We analyzed the fluorescence of the mVenus reporter tagged with unmodified ssrA or the Tat-containing sequences by flow cytometry. Only S1 showed similar levels of degradation to the control ssrA tag (**Fig. 1C**). Even though other studies showed degradation of GFP tagged with ALAA, we did not observe any degradation with S4.

While S1 was very effectively degraded, the next step was to test whether TAR RNA constructs could rescue the protein reporter by binding within the S1 tag to prevent degron recognition. Multiple WT and modified BIV TAR RNA hairpin sequences^6,22^ were cloned into a tRNA scaffold (t-TAR) for more effective folding of the hairpin and reducing RNA degradation **(Table S1)**.^23–25^ Cells co-expressing mVenus-S1 reporter and TAR constructs or no TAR as control were analyzed by flow cytometry after overnight autoinduction (**Fig. 1D**).^26^ While all cells with TAR RNA constructs recovered some fluorescence, v1, and v2 showed the most significant fluorescence increase (∼60-fold) and exhibited similar fluorescence as cells expressing untagged mVenus (85-87%). Moving forward, t-TAR-v2 was used in all subsequent experiments and will be referred to as TAR. An orthogonal induction experiment using 50 µM of isopropyl β-D-1-thiogalactopyranoside (IPTG) and 1% arabinose verified that only TAR RNA and not a binding-incompetent mutant, TAR-M1, is able to rescue cellular fluorescence (**Fig. S1A**).

In general, the degree of fluorescence activation in this reporter system should depend on the balance between the synthesis and degradation rates for the protein; the reporter system should be responsive in regimes where the change in degradation rate substantially modulates protein levels (**Fig. S1B**). However, for fluorescent protein reporters, we also considered whether chromophore maturation may be a rate-limiting step rather than synthesis. Thus, S1-tagged fluorescent proteins with different chromophore maturation times reported in FPBase^27^ were co-expressed with TAR RNA or no TAR as a control by overnight autoinduction followed by flow cytometry analysis (**Fig. 2B**). We observed that the two FPs with shorter maturation times than mVenus (13.6 min for sfGFP and 15 min for mCherry) exhibit higher background and thus substantially smaller changes in fluorescence turn-on than mVenus (17.6 min). The two FPs with longer maturation times than mVenus (25 min for EGFP and 60 min for mRFP) exhibit the opposite trend. Together, these results support that there is a maturation time threshold for the function of this reporter system (**Fig. 2C**), as FPs below a certain maturation time can form the chromophore and fluorescing before being degraded by ClpXP (**Fig. 2A**).

**Figure 2.**
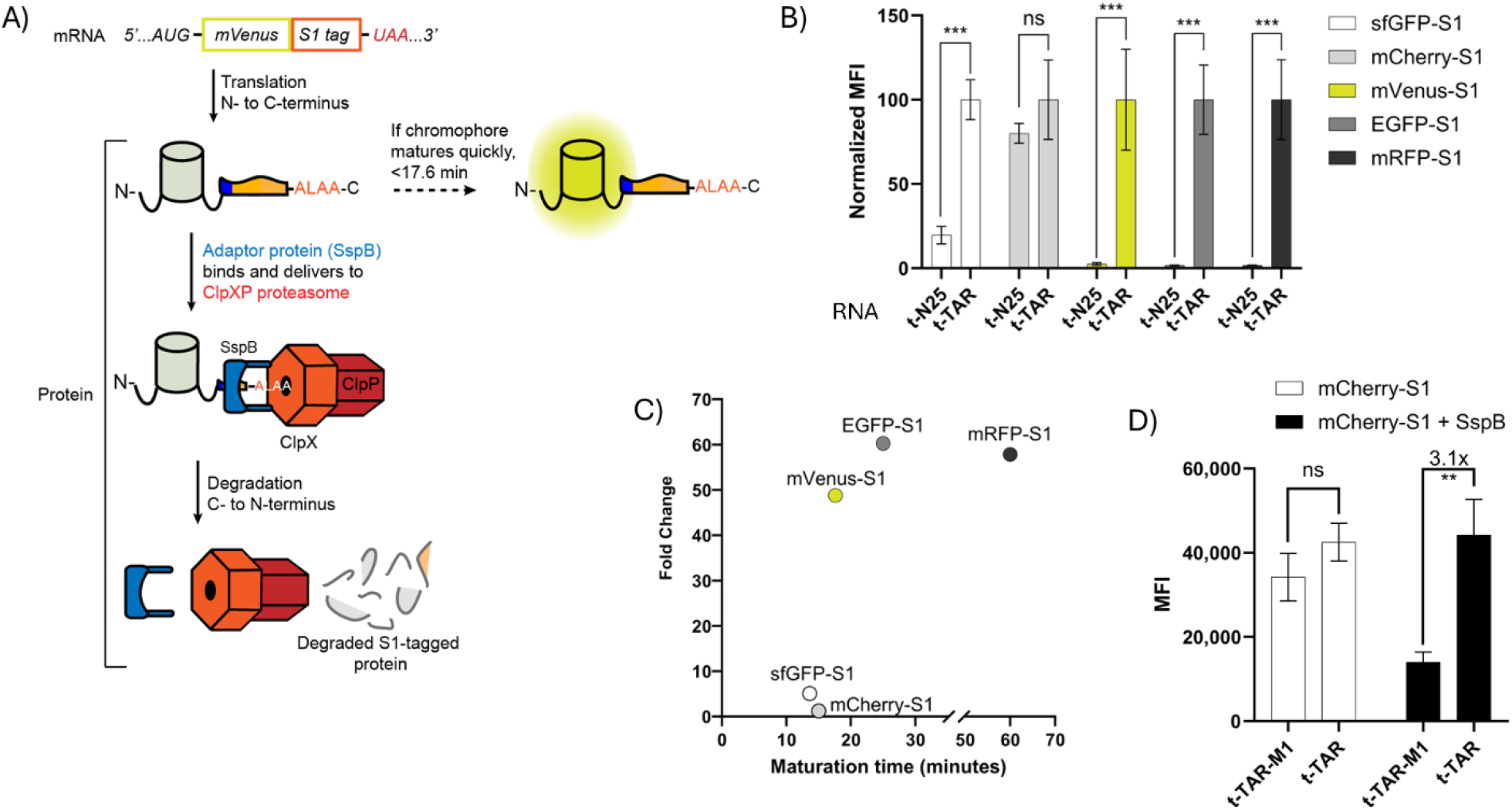
Analysis of S1-tagged fluorescent proteins with different chromophore maturation rates. (A) Scheme for how chromophore maturation and SspB levels affect background fluorescence. (B) Flow cytometry analysis of S1-tagged FPs in the presence of control (N25) or TAR RNA in tRNA scaffold (t). MFI values are normalized to the condition with TAR. (C) Graph of fluorescence fold turn-on relative to the FP maturation times. (D) Flow cytometry analysis of mCherry-S1 with and without SspB overexpression. All mean fluorescence intensity (MFI) values were analyzed for at least 100,000 cells per sample and standard deviation is shown for at least three biological replicates. TAR-M1 is a nonbinding mutant of TAR. ns, P > 0.05; *, P ≤ 0.05; **, P ≤ 0.01; ***, P ≤ 0.001.

To test this idea, we measured two additional FPs, moxGFP and mKate2 with maturation times reported in FPBase of 17.1 min and 20 min, respectively.^28^ To our initial surprise, both FPs showed higher background and little fluorescence turn-on compared to mVenus, even though their maturation times crossed the threshold (**Fig. S2**). However, the source publication for mKate2 actually reports its maturation time as “less than 20 min”,^29^ so these results suggest that its maturation time should be faster than mVenus. Furthermore, we tried to change the maturation time threshold by overexpressing the adaptor protein SspB, which was hypothesized to increase the degradation rate by ClpXP (**Fig. 2D**). As expected, increasing SspB levels in cells did reduce the fluorescence background and increased turn-on for S1-tagged mCherry, revealing this strategy to be a potential way to improve the conditional degron function.

Beyond GFP-like fluorescent proteins, we tested two other reporter proteins with the S1 tag, miniSOG^30^ and Nanoluc,^31^ both of which require binding a small molecule (chromophore or substrate) prior to their fluorescence and luminescence activity, respectively. Without increasing SspB levels, miniSOG-S1 and Nanoluc-S1 both showed significant increases in reporter signal (3.1 and 4.0-fold, respectively) in cells in the presence of TAR RNA (**Fig. S3**). Nanoluc-S1 demonstrates that the S1 conditional degron tag can be used to control enzymatic activity, not just fluorescence, and the results were readily analyzed using a plate reader.

A chief motivation to develop a conditional degron system that applies an RNA aptamer to stabilize the reporter protein is that we and other research groups have developed RNA-based biosensors to detect metabolites, signals, and regulatory small RNAs in bacteria and mammalian cells. Many of these biosensors bind an exogenous dye compound to impart the fluorescence signal for tracking and imaging in cells. The conditional degron potentially allows for a fully genetically encoded system with two expressed components, the RNA aptamer that could be converted to a biosensor and the reporter protein that imparts the fluorescence signal in place of an exogenous dye.

To simplify the system, the two components can be encoded in a single plasmid. Function of the conditional degron was compared in three dual expression plasmids (pCDF, pETDuet, and pCOLADuet), in which TAR or control RNA was encoded in one cloning site and mVenus-S1 was encoded in the other. Dual expression in pCDF yielded the highest fluorescence and fold turn-on with TAR versus control RNA (**Fig. S4A**). However, we realized that the system can be simplified even further by having both components encoded as a single transcript, driven by a single promoter. This can be accomplished by integrating the TAR RNA sequence into the 3’ untranslated region of the *mVenus-S1* mRNA, which would be translated into mVenus-S1 protein (**Fig. 3A**). The feasibility of this idea was supported by the observation that the tRNA scaffold is not needed for TAR RNA to effectively rescue fluorescence *in vivo* (**Fig. S4B**)

**Figure 3.**
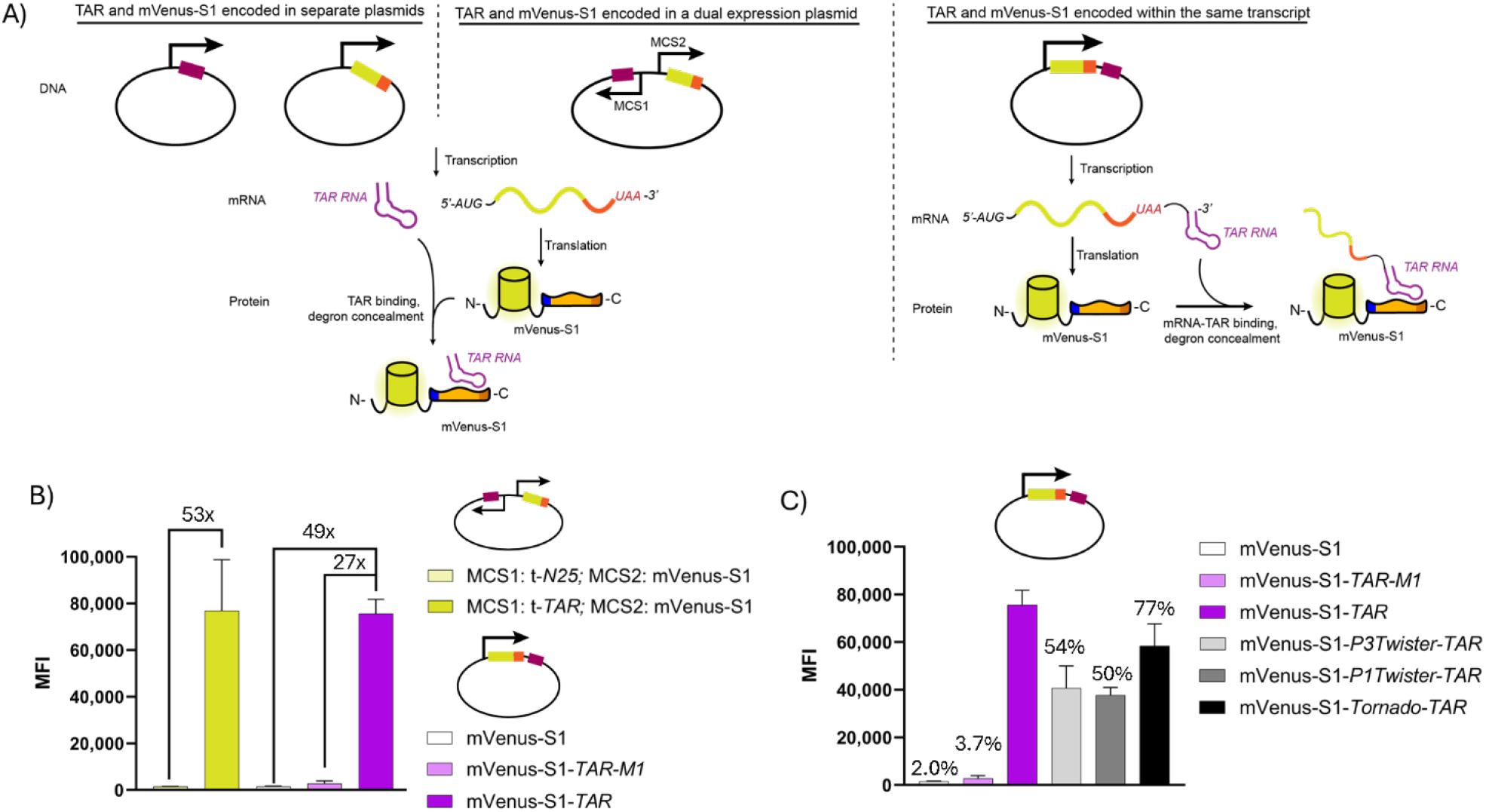
Development of a single transcript encoding both S1-tagged reporter protein and trigger TAR RNA. (A) Scheme for streamlined designs of the conditional degron system, from two plasmids to one plasmid to one transcript, where TAR is placed in the 3’ UTR. Purple is TAR RNA, yellow is mVenus, red is S1 degron tag. (B) Flow cytometry analysis to compare the one plasmid and single transcript systems. TAR-M1 is a binding incompetent mutant (**Fig. S4**). RNA constructs are in a tRNA scaffold (t) for one plasmid system only. (C) Flow cytometry analysis to compare single transcript designs, TAR retained in the full-length mRNA (TAR, TAR-M1), processed to a linear RNA by a single ribozyme (Twister), or potentially processed to a circular RNA by dual ribozymes (Tornado) (see **Fig. S5** for details). Percentages are relative to mVenus-S1-TAR. All mean fluorescence intensity (MFI) values were analyzed for at least 100,000 cells per sample and standard deviation is shown for at least three biological replicates.

We tested different mRNA constructs, in which the TAR RNA remains a part of the full-length mRNA, or the TAR RNA is designed to be processed out of the mRNA through ribozyme cleavage, with and without cyclization (**Fig. S5**). To determine the extent of fluorescence turn-on, these mRNA constructs containing TAR RNA were compared to a control mRNA containing TAR-M1, a hairpin mutant that was shown to disrupt Tat peptide binding and was found to have the lowest background for five mutants that were tested (**Fig. S4C**).^32^ Interestingly, it was found that TAR in the full-length mRNA context yielded the greatest fluorescence turn-on, comparable to dual expression of TAR and protein separately in pCDF (**Fig. 3B**). Single ribozyme-processed TAR RNAs (P3Twister U2A, P1Twister)^33^ were only half as effective, whereas the double ribozyme-processed TAR RNA (Tornado)^34^ showed improvement but still was not as fluorescent as unprocessed TAR (**Fig. 3C**).

Our initial hypothesis was that Tornado-circularized TAR RNA would be more stable *in vivo* and thus lead to greater fluorescence.^34^ While the results do support that double ribozyme-processed is more stable than single ribozyme-processed TAR, likely due to circularization, the best fluorescence turn-on was observed for the unprocessed TAR. There are two possible explanations, depending on whether the RNA or protein component of the system is limiting: either the TAR-containing mRNA is more stable than circularized TAR RNA *in vivo*, or the unprocessed mRNA produces more mVenus-S1 protein than the ribozyme-processed mRNA. In either case, we demonstrate that a single transcript encoding both TAR RNA and mVenus-S1 can be just as effective as a system with the two components encoded separately.

Moving toward sensing applications, we designed a fully genetically encodable RNA biosensor for the *E. coli* regulatory small RNA *oxyS* (**Fig. 4A**). *oxyS* is released in response to peroxide stress and has been found to regulate several proteins and mRNAs triggering temporary cell cycle arrest to allow DNA damage repair and cellular recovery.^35^ Previously, we showed that an RNA biosensor could hybridize to the small RNA *sgrS* and change folding conformation to form the fluorescent dye-binding aptamer Spinach2.^36^ However, the Spinach2-based biosensor required addition of an exogenous dye to fluoresce. More recently, other researchers showed a design of RNA biosensors to detect *survivin* mRNA and long non-coding RNA MALAT-1 that uses the original conditional degron tag in mammalian cells.^7^ Inspired by those studies, biosensors for *oxyS* RNA were designed with varied lengths of TAR truncation, sRNA targeting region, and self-complementarity region. Using the RNA folding software NUPACK ^37^ to estimate structural stability, biosensors with the most stable alternative folds that prevent TAR hairpin formation in the absence of target sRNA were selected for experimental screening. To analyze different biosensor constructs, cells were co-transformed with pET31b plasmid to express different RNA biosensors or positive and negative controls (TAR and TAR-M1, respectively) and pCDF plasmid to dually express the reporter mVenus-S1 and the sRNA (either the target *oxyS* or non-target *pinT*). Of the five sensor designs that were screened, several appeared to display higher fluorescence activation in the presence of *oxyS*, although there was high variance in the fluorescence signal (**Fig. S6B**). Upon retesting, we observed ∼15-fold increase in fluorescence for one of the biosensors, TARoxyS-A6, in the presence of the *oxyS* RNA target relative to the non-target (**Fig. 4B**). In contrast, expression of the sRNAs did not significantly affect TAR or TAR-M1, which showed constitutively high and low fluorescence, as expected.

**Figure 4.**
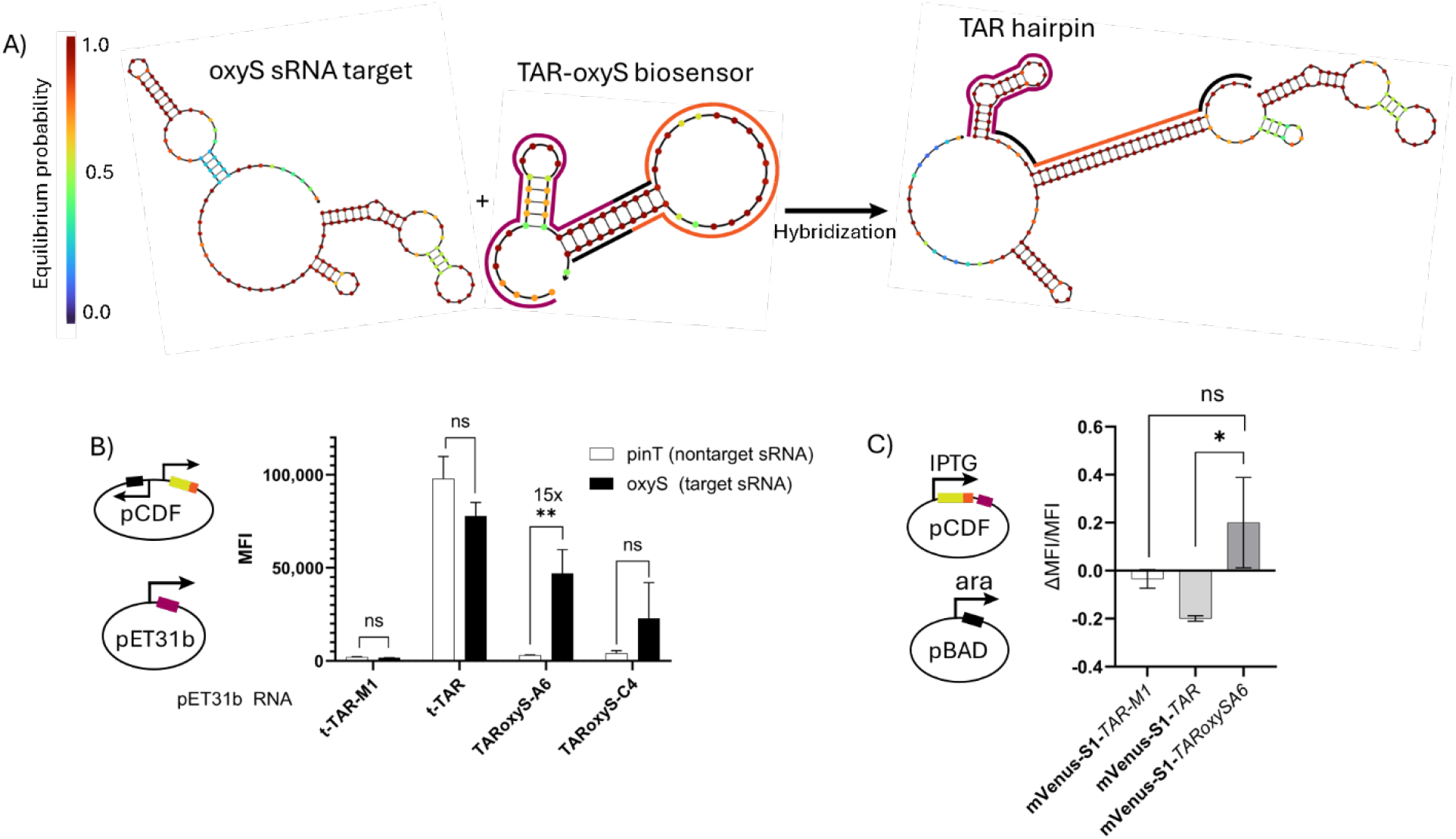
Proof-of-concept application of the conditional degron to construct a TAR-based sRNA biosensor. (A) NUPACK predicted secondary structure for *oxyS* sRNA, TAR-oxyS biosensor, and predicted hybridization between the sRNA and biosensor to form the TAR hairpin. For the TAR-oxyS biosensor, the sequence for the TAR RNA hairpin is highlighted in maroon, self-complementary regions are highlighted in black, sRNA-targeting sequence is highlighted in orange and is the reverse complement to the sRNA target. (B) Flow cytometry analysis of two biosensor designs, A6 and C4, in pET31b in the presence of target (*oxyS*) or non-target (*pinT*) sRNA (also see **Fig. S6**). TAR and TAR-M1 in a tRNA scaffold (t) are additional controls. MFI values were analyzed for at least 100,000 cells per sample and the standard error of the mean is shown for 4-8 biological replicates. (C) Flow cytometry analysis of the one plasmid conditional degron system using the TAR-oxyS-A6 biosensor with and without induction of *oxyS* sRNA expression by arabinose (also see **Fig. S6**). MFI values were analyzed for least 100,000 cells per sample and standard deviation is shown for three biological replicates. For each biological replicate, ΔMFI/MFI values were calculated by subtracting MFI_+IPTG/-Ara_ from MFI_+IPTG/+Ara_ and dividing by MFI_+IPTG/-Ara_

As proof-of-concept for the single transcript biosensor, we integrated the TARoxyS-A6 biosensor or positive and negative controls (TAR and TAR-M1, respectively) into the 3’ untranslated region of the *mVenus-S1* mRNA (**Supplementary Table 3**). Cells were co-transformed with the pCDF plasmid expressing this single transcript biosensor under IPTG control and the pBAD plasmid expressing *oxyS* sRNA under arabinose control. As expected, little to no cellular fluorescence is observed for single transcript containing TAR-M1 under any conditions, and fluorescence requires IPTG induction of single transcripts containing TAR or TARoxyS-A6 (**Fig. S6C**). Cellular fluorescence for bioreplicates were analyzed after 3 h without or with arabinose induction of *oxyS* sRNA (**Fig. 4C**). Single transcript containing TAR-M1 exhibited no change from very low fluorescence, consistent with the complete absence of a binding-competent TAR sequence. Interestingly, single transcript containing TAR exhibited a slight fluorescence decrease, which could be attributed either to arabinose induction competing slightly with expression or some effect of *oxyS* sRNA. In contrast, the single transcript containing TARoxyS-A6 exhibited a slight fluorescence increase, which suggests that it can detect *oxyS* sRNA expression.

To our knowledge, this is the very first example of a novel biosensor design that enables expressing both protein and RNA components from a single transcript. The TARoxyS-A6 biosensor displayed higher background fluorescence (without *oxyS*) in the single transcript than in the dual expression system, though. This result indicates that the biosensor sequence is more likely to form the TAR hairpin instead of the alternative structure modeled by NUPACK when in the full-length mRNA than in a standalone RNA transcript. Optimizing the mRNA sequence context or applying the Tornado system could improve the fluorescence response for the biosensor.

## CONCLUSIONS

In this study, we developed a conditional degron system for use in bacteria in which an FP reporter, mVenus-S1, is stabilized only in the presence of an RNA, TAR. The functional design takes advantage of the bipartite nature of the ssrA C-terminal tag, so that TAR RNA binding to the inserted Tat peptide sequence blocks degron recognition to rescue protein activity. Conversely, overexpressing the SspB adapter protein can decrease background fluorescence to improve the fold response. We envision that this mVenus-S1/TAR system could be used as a reporter for studying transcriptional activity from promoters and/or tracking transcripts tagged with the TAR hairpin, which was shown to fold efficiently with or without a scaffold and in an mRNA. Because mVenus-S1 protein is degraded when not bound to the RNA, the conditional degron tag should have lower background fluorescence than existing tags that use FP-RNA-binding protein fusions.^38^ The TAR hairpin is similar in size to MS2 and PP7 RNA hairpins (20-30 nucleotides).^39^

In addition, we showed that chromophore maturation time correlates with the fold response of the conditional degron, with slow maturing FPs having the highest fold response. The responsiveness cutoff is in the range of 17.1-17.6 min but can be shifted to shorter maturation times by increasing SspB levels. In fact, our experiments indicate that mKate2, which was reported to have a maturation rate of <20 min, appears to have a faster maturation rate than mVenus (< 17.6 min). Furthermore, other types of protein reporters can be applied for RNA sensing, such as miniSOG and Nanoluc. Taken together, these results demonstrate that the conditional degron system has potential for controlling protein activity in bacteria as triggered by RNA inputs.

Finally, we designed a functional biosensor for *oxyS* sRNA and integrated it into a single transcript in which the FP and RNA-based biosensor are encoded on the same mRNA. While our experiments only provide proof of principle, this is the first demonstration of a streamlined system that is fully genetically encodable, does not require an exogenous fluorophore, and permits both components to be coordinated by expression from the same promoter. Our results support that some of the Tornado-processed TAR were being circularized by the native RtcB RNA ligase,^40^ as they showed stronger fluorescence than the single ribozyme constructs that yield linear TAR. Ongoing work focuses on improving these single transcript biosensors and applying them to sense novel targets.

