## Supplementary material for "A Conditional Degron Tag for RNA Sensing in Bacteria": Methods and SI

Table of Contents

| Figure label | Description | Pages |
|---|---|---|
| **Figure S1** | Orthogonal induction of mVenus-S1 protein and TAR RNA | S2 |
| **Figure S2** | Analysis of S1-tagged fluorescent protein degradation and rescue by t-TAR RNA | S3 |
| **Figure S3** | Analysis of additional S1-tagged protein degradation and rescue by t-TAR RNA | S4 |
| **Figure S4** | Optimization of how TAR RNA and mVenus-S1 protein are encoded. | S5 |
| **Figure S5** | Schematics of binding of TAR when encoded in 3'UTR of reporter mRNA | S6 |
| **Figure S6** | Development of a TAR-based oxyS sRNA biosensor | S7 |
| **Table S1** | RNA sequences used in this work | S8 |
| **Table S2** | Plasmids used in this work | S9-S10 |
| **Table S3** | DNA sequences related to this work | S11-S12 |
| **Materials and Methods** | | S13 |
| **References** | | S15 |

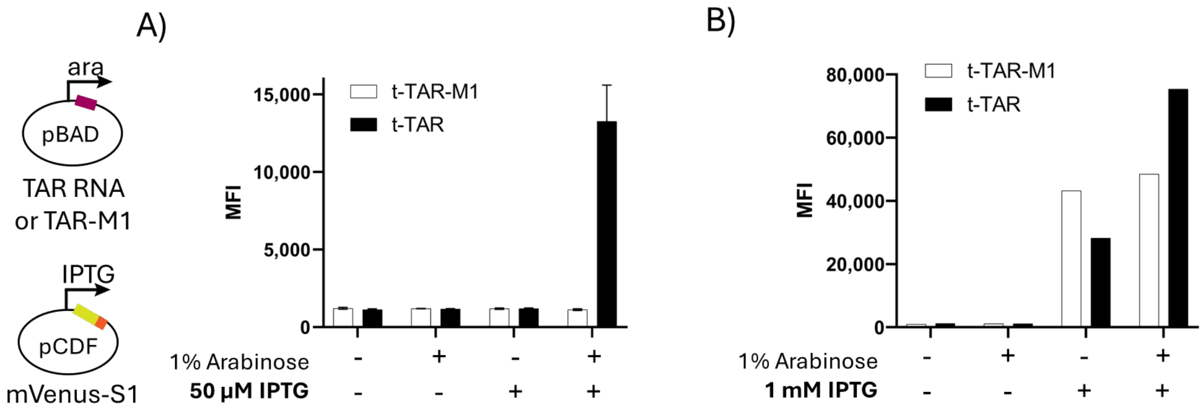

**Figure S1: Orthogonal induction of mVenus-S1 protein and TAR RNA.** Flow cytometry analysis of mVenus-S1 with and without induction of TAR RNA expression by arabinose. Mean fluorescence intensity (MFI) values were analyzed for at least 100,000 cells per sample. A) Standard deviation is shown for three biological replicates treated with and without 50 µM IPTG. B) A single biological sample per RNA treated with and without 1 mM IPTG.

An unusually low concentration of 50 µM IPTG was required to slow the expression rate of mVenus-S1 to not outpace the degradation rate while allowing measurable MFI within four hours of induction. Typical IPTG concentrations (0.1-1 mM IPTG) resulted in an expression rate faster than mVenus-S1 could be degraded resulting in a "false positive" in fluorescence rescue. The "+IPTG/-Ara" samples display this effect the most clearly.

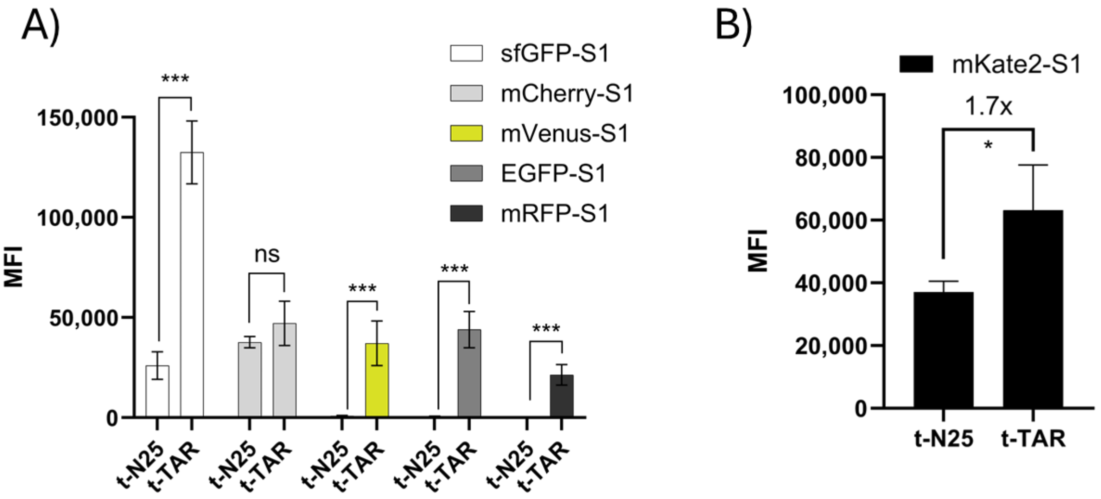

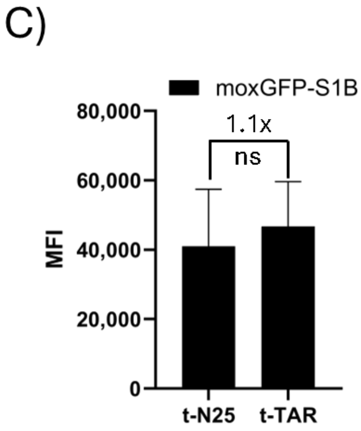

D)

| FP | Maturation time (min) | Fold change |
|---|---|---|
| sfGFP-S1 | 13.6 | 5.1 |
| mCherry-S1 | 15 | 1.2 (ns) |
| *moxGFP-S1* | *17.1* | *1.1 (ns)* |
| *mKate2-S1* | *<20* | *1.7* |
| mVenus-S1 | 17.6 | 49 |
| EGFP-S1 | 25 | 60 |
| mRFP-S1 | 60 | 58 |

**Figure S2: Analysis of S1-tagged fluorescent protein degradation and rescue by TAR RNA.** Flow cytometry analysis of S1-tagged FPs in the presence of control (N25) or TAR RNA in tRNA scaffold (t). Mean fluorescence intensity (MFI) values were analyzed for least 100,000 cells per sample and standard deviation is shown for at least three biological replicates. A) S1-tagged FP MFI values before normalization. B) Flow cytometry analysis of mKate2-S1. C) Flow cytometry analysis of moxGFP-S1. A much lower blue laser gain of 320 (rather than normal 540) was required to measure the full cell fluorescence histogram. D) Compiled reported maturation times (FPBase)[1] and observed fold change from t-N25 to t-TAR RNA. moxGFP-S1 and mKate2-S1 are italicized, because they are excluded from **Fig 2B** and **Fig. S2A.**

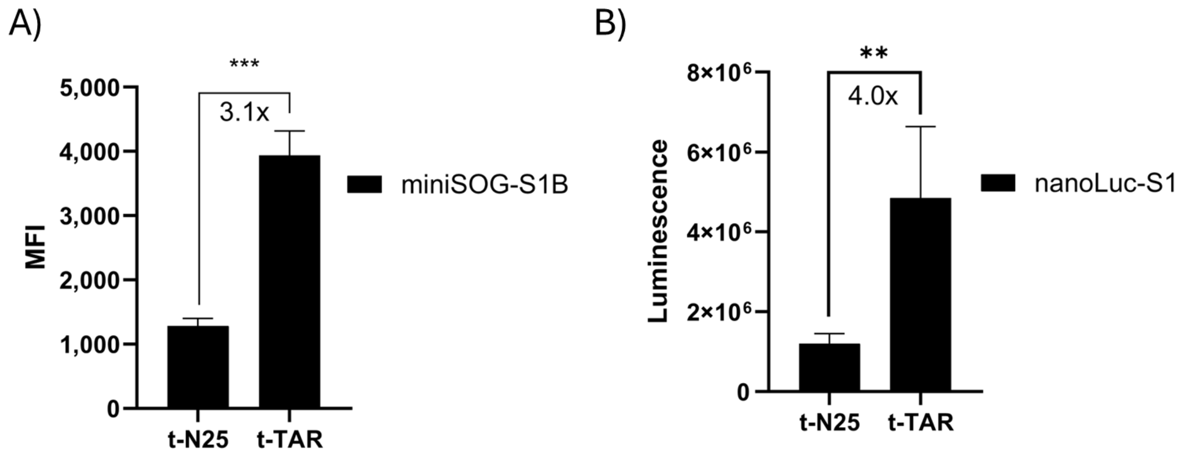

**Figure S3: Analysis of additional S1-tagged protein degradation and rescue by t-TAR RNA.** A) Flow cytometry analysis of miniSOG-S1 in the presence of control (N25) or TAR RNA in tRNA scaffold (t). Mean fluorescence intensity (MFI) values were analyzed for least 100,000 cells per sample and standard deviation is shown for at least three biological replicates. Addition of riboflavin to media during expression did not affect fluorescence in either untagged or tagged miniSOG (data not shown). B) Plate reader analysis of autoinduction cultures expressing nanoLuc-S1 in the presence of control (N25) or TAR RNA in tRNA scaffold (t). Measurements were taken two minutes after addition of chemiluminescent substrate. Plate reader luminescence values shown are the average with a standard deviation of three biological replicates.

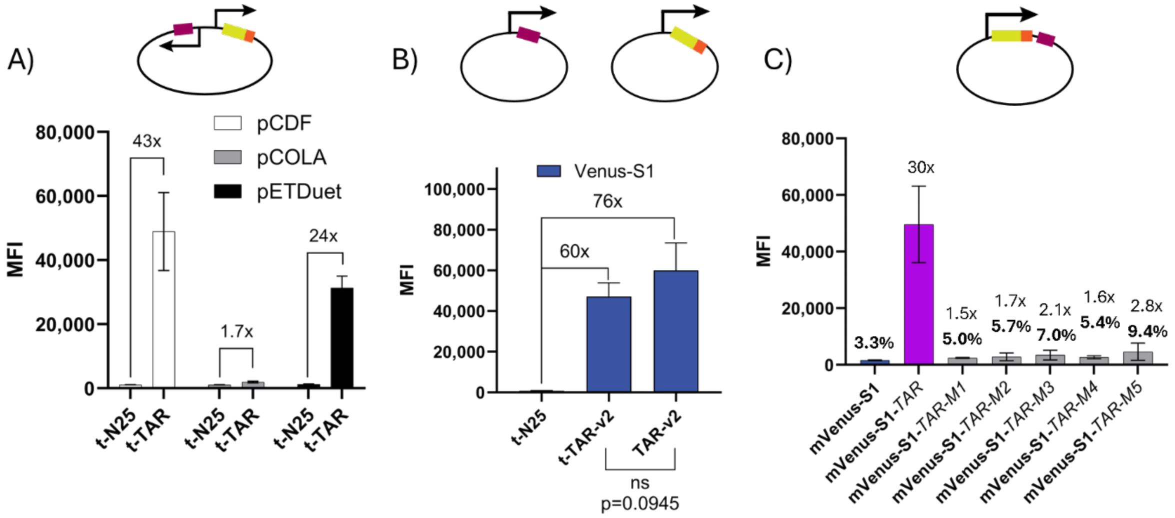

**Figure S4: Optimization of how TAR RNA and mVenus-S1 protein are encoded.** Mean fluorescence intensity (MFI) values were analyzed for least 100,000 cells per sample and standard deviation is shown for at least three biological replicates A) Flow cytometry analysis comparing dual expression plasmids. Genes were cloned to be divergent on opposite strands following for more consistent expression.[2] B) Flow cytometry analysis of mVenus-S1 comparing TAR RNA with and without a tRNA scaffold (t) when RNA is encoded in a separate pET31b plasmid. C) Flow cytometry analysis of designed nonbinding TAR mutants in the context of the 3' UTR of mVenus-S1 mRNA.

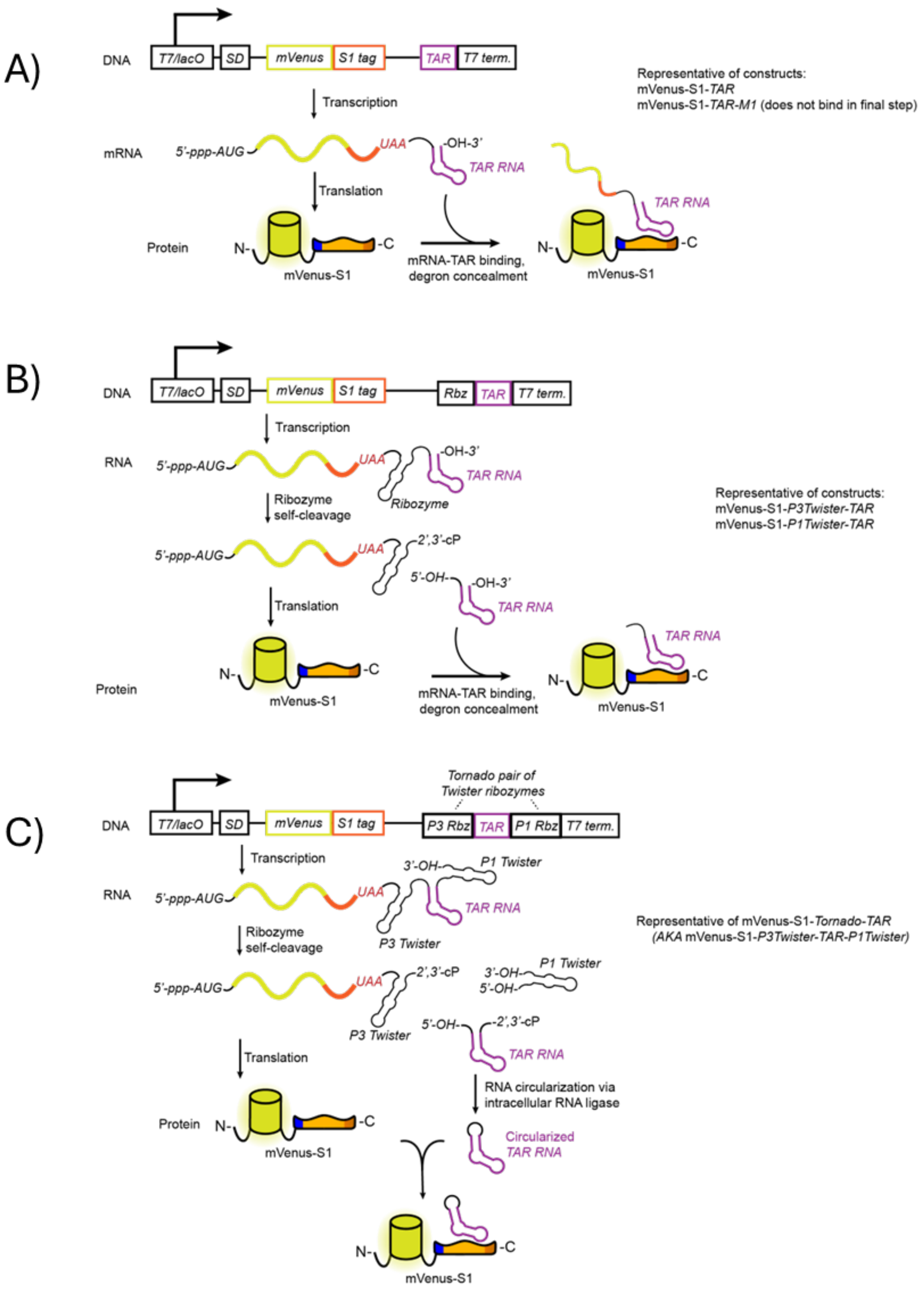

**Figure S5: Schematics of proposed binding of TAR when encoded in 3'UTR of reporter mRNA.** A) Constructs designed for no cleavage of TAR from mRNA, B) designed for TAR to be cleaved from mRNA via a single ribozyme but not circularized,[3] and C) and designed for TAR to be cleaved from mRNA via a ribozyme pair and subsequently circularized by an endogenous RtcB RNA ligase.[4,5]

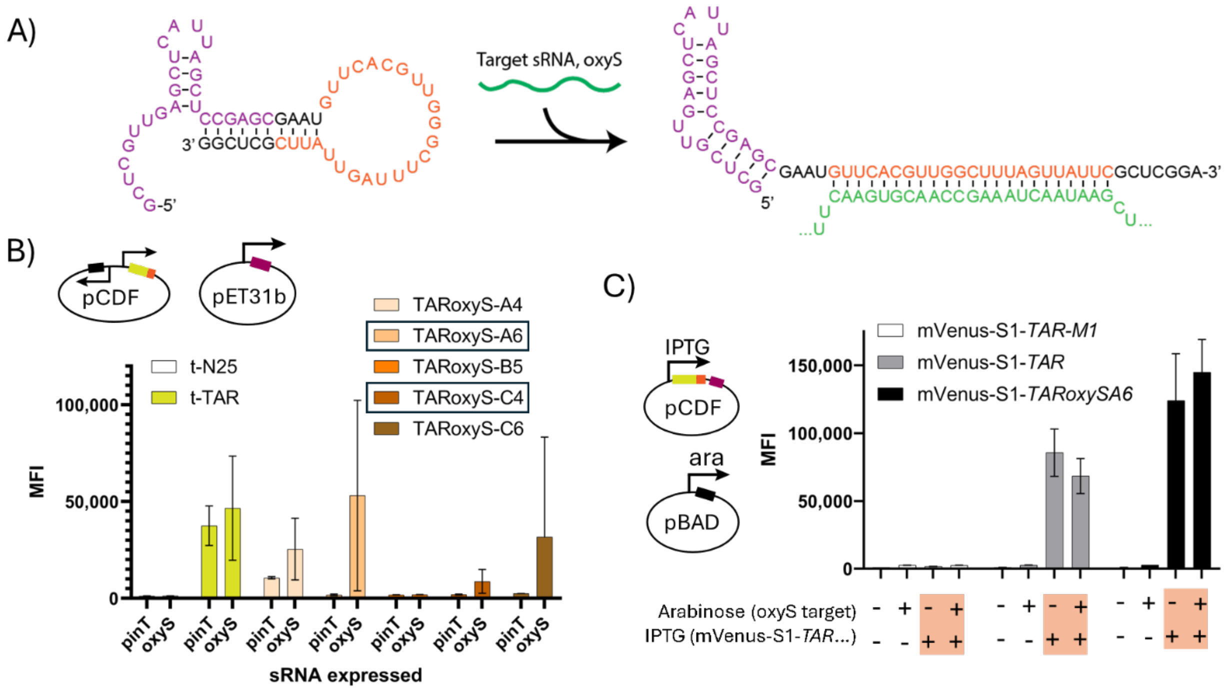

**Figure S6: Development of a TAR-based** oxyS **sRNA biosensor.** A) Scheme showing the predicted secondary structure of the TARoxyS-A6 sensor before and after target hybridization. B) Initial screening via flow cytometry analysis of five biosensor designs in pET31b in the presence of target (*oxyS*) or non-target (*pinT*) sRNA. TAR and TAR-M1 in a tRNA scaffold (t) are additional controls. MFI values were analyzed for at least 100,000 cells per sample and standard error of the mean is shown for three biological replicates. C) Flow cytometry analysis of the one plasmid conditional degron system using the TARoxyS-A6 biosensor with and without induction of *oxyS* sRNA expression by arabinose. Mean fluorescence intensity (MFI) values were analyzed for at least 100,000 cells per sample and standard deviation is shown for at least three biological replicates.

The variance in MFI between biological replicates concealed biosensing that occurred from "+IPTG/-Ara" to "+IPTG/+Ara". Because the same biological sample was split for different induction conditions, we paired the subcultures to show the change in MFI when both IPTG and arabinose were added. The ΔMFI/MFI values shown in **Fig. 4C** were more indicative of the MFI increase observed upon oxyS sRNA induction (via arabinose) for TARoxyS-A6 in the mVenus-S1 mRNA context.

**Table S1: RNA sequences used in this work.** tRNA scaffold sequences are shown in bold. TAR hairpin bases are shown in purple. Nucleobases that are changed from its wildtype or parent RNA are shown underlined. oxyS-targeting regions are shown in green.

| RNA ID | RNA Sequence (5' to 3') |
|---|---|
| t-N25 | **GCCCGGAUAGCUCAGUCGGUAGAGCAGCGGCCG**AUAAAAUCACCAGUACCCAAGACCA**CGGCCGCGGGUCCAGGGUUCAAGUCCCUGUUCGGGCGCCA** |
| t-TAR-wt | **GCCCGGAUAGCUCAGUCGGUAGAGCAGCGGCCG**GGCUCGUGUAGCUCAUUAGCUCCGAGCC**CGGCCGCGGGUCCAGGGUUCAAGUCCCUGUUCGGGCGCCA** |
| t-TAR-v1 | **GCCCGGAUAGCUCAGUCGGUAGAGCAGCGGCCG**GGCUCGUCUGAGCUCAUUAGCUCCGAGCC**CGGCCGCGGGUCCAGGGUUCAAGUCCCUGUUCGGGCGCCA** |
| t-TAR-v2 | **GCCCGGAUAGCUCAGUCGGUAGAGCAGCGGCCG**GGCUCGUUGAGCUCAUUAGCUCCGAGCC**CGGCCGCGGGUCCAGGGUUCAAGUCCCUGUUCGGGCGCCA** |
| t-TAR-v3 | **GCCCGGAUAGCUCAGUCGGUAGAGCAGCGGCCG**GGCUCGUUGAGCUCUGGGAAGCUCCGAGCC**CGGCCGCGGGUCCAGGGUUCAAGUCCCUGUUCGGGCGCCA** |
| TAR-v2 ("TAR") | GGCUCGUUGAGCUCAUUAGCUCCGAGCC |
| TAR-M1 (G11C C20G) | GGCUCGUUGACCUCAUUAGGUCCGAGCC |
| TAR-M2 (C20U) | GGCUCGUUGAGCUCAUUAGUUCCGAGCC |
| TAR-M3 (C12G G19C) | GGCUCGUUGAGGUCAUUACCUCCGAGCC |
| TAR-M4 (G19C) | GGCUCGUUGAGCUCAUUACCUCCGAGCC |
| TAR-M5 (G19C, C20U, U21A, C22U) | GGCUCGUUGAGCUCAUUACUAUCGAGCC |
| t-TAR-M1 (G11C C20G) | **GCCCGGAUAGCUCAGUCGGUAGAGCAGCGGCCG**GGCUCGUUGACCUCAUUAGGUCCGAGCC**CGGCCGCGGGUCCAGGGUUCAAGUCCCUGUUCGGGCGCCA** |
| pinT sRNA | AGUAACGGAUUACUUUGUGGUGUAGCGUAACGGUAAUUGUCCUCCUCAUAUUUGCGGCAGCGUAGUCUGCCGCUUUUUUU |
| oxyS sRNA target | GAAACGGAGCGGCACCUCUUUUAACCCUUGAAGUCACUGCCCGUUUCGAGAGUUUCUCAACUCgaauaacuaaagccaacgugaacUUUUGCGGAUCUCCAGGAUCCGCUUUUUUUUGCCAUAAAAAA |
| TAR-oxyS-A4 | GCUCGUUGAGCUCAUUAGCUCCGAGCGAAUAAGUUCACGUUGGCUUUAGUUAUUCGCUC |
| TAR-oxyS-A6 | GCUCGUUGAGCUCAUUAGCUCCGAGCGAAUGUUCACGUUGGCUUUAGUUAUUCGCUCGG |
| TAR-oxyS-B5 | UUGAGCUCAUUAGCUCGAAUAGUUCACGUUGGCUUUAGUUAUUCGAGCU |
| TAR-oxyS-C4 | UUGAGCUCAUUAGCUCACUAAAGUUGGCUUUAGUGAGC |
| TAR-oxyS-C6 | UUGAGCUCAUUAGCUCACUAGUUGGCUUUAGUGAGCUA |

**Table S2: Plasmids used in this work.** Detailed plasmid maps are supplied as a ZIP file, or at the following link https://benchling.com/tjsimons763/f_/cmuGhQOe-s1-tag-manuscript-file-copies/

| Plasmid backbone | MCS1 | MCS2 | Used in figures | Antibiotic resistance |
|---|---|---|---|---|
| pCDF | Empty | mVenus | **Figs. 1C-D,** | Spectinomycin |
| | | mVenus-ssrA | **Fig. 1C** | |
| | | mVenus-S1 | **Figs. 1C-D, 2B-C, 3B-C, S1, S2A, S2D, S4C** | |
| | | mVenus-S2 | **Fig. 1C** | |
| | | mVenus-S3 | **Fig. 1C** | |
| | | mVenus-S4 | **Fig. 1C** | |
| pET31b | N/A | t-N25 | **Figs. 1D, 2B-D, S2A-D, S3, S4B, S6B** | Carbenicillin |
| | | t-TAR-wt | **Fig. 1D** | |
| | | t-TAR-v1 | **Fig. 1D** | |
| | | t-TAR-v2 (“t-TAR” and “TAR”) | **Figs. 1D, 2B-D, S2A-D, S3, S4B, S6B** | |
| | | t-TAR-v3 | **Fig. 1D** | |
| pCDF | Empty | sfGFP-S1 | **Figs. 2B-C, S2A, S2D** | Spectinomycin |
| | | mCherry-S1 | **Figs. 2B-D, S2A, S2D** | |
| | | EGFP-S1 | **Figs. 2B-C, S2A, S2D** | |
| | | mRFP-S1 | **Figs. 2B-C, S2A, S2D** | |
| pCDF | SspB | mCherry-S1 | **Fig. 2D** | |
| | Empty | mVenus-S1-TAR-M1 (G11C C20G) | **Figs. 3B-C, 4C, S4C, S5A, S6C** | |
| | | mVenus-S1-TAR | **Figs. 3B-C, 4C, S4C, S5A, S6C** | |
| | | mVenus-S1-P3TwisterU2A-TAR | **Figs. 3C, S5B** | |
| | | mVenus-S1-P1TwisterTAR | **Figs. 3C, S5B** | |
| | | mVenus-S1-Tornado-TAR | **Figs. 3C, S5C** | |
| | tN25 | mVenus-S1 | **Figs. 3B, S4A** | |
| | tTAR | | **Figs. 3B, S4A** | |
| | pinT sRNA | | **Fig. 4B** | |

| | | | | |
|---|---|---|---|---|
| | oxyS sRNA | | **Fig. 4B** | |
| pET31b | N/A | t-TAR-M1 | **Fig. 4B** | Carbenicillin |
| | | TARoxyS-A4 biosensor | **Fig. S6B** | |
| | | TARoxyS-A6 biosensor | **Figs. 4B, S6A-B** | |
| | | TARoxyS-B5 biosensor | **Fig. S6B** | |
| | | TARoxyS-C4 biosensor | **Figs. 4B, S6B** | |
| | | TARoxyS-C6 biosensor | **Fig. S6B** | |
| pCDF | Empty | mVenus-S1-TARoxyS-A6 (single transcript biosensor) | **Fig. 4C, S6C** | Spectinomycin |
| pBAD | N/A | oxyS sRNA target | **Fig. 4C** | Carbenicillin |
| | | t-TAR | **Fig. S1** | |
| | | t-TAR-M1 | **Fig. S1** | |
| pCDF | Empty | moxGFP-S1 | **Figs. S2C-D** | Spectinomycin |
| | | mKate2-S1 | **Figs. S2B, S2D** | |
| | | miniSOG-S1 | **Fig. S3A** | |
| | | nanoLuc-S1 | **Fig. S3B** | |
| pCOLA | tN25 | mVenus-S1 | **Fig. S4A** | Kanamycin |
| | tTAR | | **Fig. S4A** | |
| pETDuet | tN25 | | **Fig. S4A** | Carbenicillin |
| | tTAR | | **Fig. S4A** | |
| pET31b | N/A | TAR-v2 | **Fig. S4B** | |
| pCDF | Empty | mVenus-S1-TAR-M2 (C20U) | **Fig. S4C** | Spectinomycin |
| | | mVenus-S1-TAR-M3 (C12G G19C) | **Fig. S4C** | |
| | | mVenus-S1-TAR-M4 (G19C) | **Fig. S4C** | |
| | | mVenus-S1-TAR-M5 (G19C, C20U, U21A, C22U) | **Fig. S4C** | |

**Table S3: DNA sequences related to this work.**

| ID | Use | DNA Sequence (5' to 3') | Description |
|---|---|---|---|
| Primer 1 | Amplification of pCDF backbone with S1 tag to swap tagged protein | GGCGGAAGATCCGGTGGT | Forward primer located on linker before S1 tag |
| Primer 2 | | atgtatatctccttcttatacttaactaatatactaagatggggaattgttatccg | pCDF Reverse primer, located immediately before start codon |
| mVenus-S1-TAR | Encoding TAR into 3' UTR of mVenus-S1. Inserted ribozymes cleave TAR from the mRNA | ...ggcggaagatccggtggtggtGCTGCAAATGACGAAAATTATcgtccccgtggtactcgtggtaaaggtcgccgtattcgccgtGCTTTGGCGGCCTAAgcagatctcaattggatatcggccggccacgcgatcgctgacgtcggtaccctcgagtctGGCTCGTTGAGCTCATTAGCTCCGAGCC-3' | DNA sequences are encoded exactly after the stop codon of the tagged protein. Nonbold bases are linker regions, orange bases depict S1-tag, purple text represents TAR-v2, **bold typeface represents ribozyme bases,** underlined bases represent point mutations or TARoxyS sensor |
| mVenus-S1-P3TwisterU2A-TAR | | ...ggcggaagatccggtggtggtGCTGCAAATGACGAAAATTATcgtccccgtggtactcgtggtaaaggtcgccgtattcgccgtGCTTTGGCGGCCTAAgcagatctcaattggatatcggccggccacgcgatcgctgacgtcggtaccctcgagtct**tggccgcactcgccggtcccaagcccggataaaatgggagggggcgggaaaccgcctaaccatgccgagt**GGCTCGTTGAGCTCATTAGCTCCGAGCC-3' | |
| mVenus-S1-P1TwisterTAR | | ...ggcggaagatccggtggtggtGCTGCAAATGACGAAAATTATcgtccccgtggtactcgtggtaaaggtcgccgtattcgccgtGCTTTGGCGGCCTAAgcagatctcaattggatatcggccggccacgcgatcgctgacgtcggtaccctcgagtct**tcggcgtggactgtagaacactgccaatgccggtcccaagcccggataaaagtggagggtacagtccacgc**GGCTCGTTGAGCTCATTAGCTCCGAGCC-3' | |
| mVenus-S1-Tornado-TAR | | ...ggcggaagatccggtggtggtGCTGCAAATGACGAAAATTATcgtccccgtggtactcgtggtaaaggtcgccgtattcgccgtGCTTTGGCGGCCTAAgcagatctcaattggatatcggccggccacgcgatcgctgacgtcggtaccctcgagtct**ggccgcactcgccggtcccaagcccggataaaatgggagggggcgggaaaccgcctaaccatgccgagt**gcggccgcGGCTCGTTGAGCTCATTAGCTCCGAGCCgtggccgcgg**tcggcgtggactgtagaacactgccaatgccggtcccaagcccggataaaagtggagggtacagtccacgc**-3' | |
| mVenus-S1-TAR-M1 (G11C C20G) | Nonbinidng mutants of mVenus-S1-TAR | ...ggcggaagatccggtggtggtGCTGCAAATGACGAAAATTATcgtccccgtggtactcgtggtaaaggtcgccgtattcgccgtGCTTTGGCGGCCTAAgcagatctcaattggatatcggccggccacgcgatcgctg | |

| | | |
|---|---|---|
| | | acgtcggtaccctcgagtctGGCTCGTTGACCTCATTAGGTCCGAGCC-3' |
| mVenus-S1-TAR-M2 (C20U) | | ...ggcggaagatccggtggtggtGCTGCAAATGACGAAAATTATcgtccccgtggtactcgtggtaaaggtcgccgtattcgccgtGCTTTGGCGGCCTAAgcagatctcaattggatatcggccggccacgcgatcgctgacgtcggtaccctcgagtctGGCTCGTTGAGCTCATTAGTTCCGAGCC-3' |
| mVenus-S1-TAR-M3 (C12G G19C) | | ...ggcggaagatccggtggtggtGCTGCAAATGACGAAAATTATcgtccccgtggtactcgtggtaaaggtcgccgtattcgccgtGCTTTGGCGGCCTAAgcagatctcaattggatatcggccggccacgcgatcgctgacgtcggtaccctcgagtctGGCTCGTTGAGGTCATTACCTCCGAGCC-3' |
| mVenus-S1-TAR-M4 (G19C) | | ...ggcggaagatccggtggtggtGCTGCAAATGACGAAAATTATcgtccccgtggtactcgtggtaaaggtcgccgtattcgccgtGCTTTGGCGGCCTAAgcagatctcaattggatatcggccggccacgcgatcgctgacgtcggtaccctcgagtctGGCTCGTTGAGCTCATTACCTCCGAGCC-3' |
| mVenus-S1-TAR-M5 (G19C, C20U, U21A, C22U) | | ...ggcggaagatccggtggtggtGCTGCAAATGACGAAAATTATcgtccccgtggtactcgtggtaaaggtcgccgtattcgccgtGCTTTGGCGGCCTAAgcagatctcaattggatatcggccggccacgcgatcgctgacgtcggtaccctcgagtctGGCTCGTTGAGCTCATTACTATCGAGCC-3' |
| mVenus-S1-TARoxyS-A6 (single transcript biosensor) | Functional TAR-based oxyS sRNA biosensor | ...ggcggaagatccggtggtggtGCTGCAAATGACGAAAATTATcgtccccgtggtactcgtggtaaaggtcgccgtattcgccgtGCTTTGGCGGCCTAAgcagatctcaattggatatcggccggccacgcgatcgctgacgtcggtaccctcgagtctggtGCTCGTTGAGCTCATTAGCTCCGAGCgaatgttcacgttggctttagttattcGCTCGG-3' |

### MATERIALS AND METHODS

#### General Transformation Protocol

A 30 µL aliquot of chemically competent frozen BL21 Star (DE3) or XL1-Blue *E. coli* (Berkeley Macrolab) was combined with ~100 ng of plasmid. For co-transformations with two plasmids, ~100 ng of each plasmid was added to the frozen cell aliquot. The aliquot was allowed to thaw on ice for 30 min. The cells were then heat shocked at 42 °C for 60 seconds on a heat block and immediately placed on ice for 5 minutes. Afterward, 500 µL of LB media were added, and the cells recovered for 45 min in an incubator set to 37°C shaking at 250 rpm. After recovering, 40 µL of cells in media were plated on LB agar plates containing 50 µg/mL of the antibiotic carbenicillin, spectinomycin, or kanamycin, depending on the plasmid transformed into the cells. In the case of co-transformations, 100 µL of the 500 µL of recovered cells in media were plated on LB agar plates containing 50 µg/mL of carbenicillin and spectinomycin, respectively. The plates were incubated at 37 °C for 16–18 h.

#### General Plasmid Construction Protocol

All plasmids used in this work can be found in Supplementary Table 2 and were assembled using either Gibson DNA assembly or PCR mutagenesis followed by blunt end ligation. The Gibson assembly of protein genes into the desired plasmid was achieved using primer overhangs of 35 base pairs complementary to the plasmid backbone. PCR mutagenesis uses DNA mutations encoded in overhangs, but overlapping complementary DNA sequences are not required for blunt-end ligations. Primers were designed manually using the NEB Tm calculator, and primers for cloning a different S1-tagged protein can be found in Supplementary Table 3.

PCRs of plasmid backbones were performed using Q5 High-Fidelity DNA Polymerase enzyme (New England Biolabs) in a 50 µL reaction volume (28.5 µL sterile water, 10 µL 5X Q5 reaction buffer, 2.5 µL each of 10 µM forward and reverse primers, 1 µL template DNA at a concentration of 1 ng/µL, 5 µL 2.5 mM dNTPs, and 0.5 µL of polymerase). Thermocycling conditions were 98°C initial denaturation for 30 s, followed by 34 cycles at 98 °C for 30 s for denaturing, annealing for 30 s (specific annealing temperatures were calculated for each primer pair using the NEB Tm calculator), and extension at 72 °C for 210 s. Thermocycling was completed with a final extension at 72 °C for 5 min.

PCRs of protein, RNA constructs, and target sRNA genes were performed using Phusion DNA polymerase (Berkeley Macrolab) in a 50 µL reaction volume (28.5 µL sterile water, 10 µL 5X Phusion HF buffer, 2.5 µL each of 10 µM forward and reverse primers, 1 µL template DNA at a concentration of 1 ng/µL, 5 µL 2.5 mM dNTPs, and 0.5 µL of polymerase). Thermocycling conditions were 98°C initial denaturation for 30 s, followed by 34 cycles at 98 °C for 30 s for denaturing, annealing for 30 s (specific annealing temperatures were calculated for each primer pair using the NEB Tm calculator), and extension at 72 °C for 45 s. Thermocycling was completed with a final extension at 72 °C for 5 min.

Agarose gels were used to verify DNA amplification at the correct size using 5 µL of the PCR product. The remaining 45 µL of DNA product was treated with 1 µL of Dpn1 digest enzyme (New

England Biolabs) and incubated at 37 °C for 1 h. The DNA was then purified using Qiagen PCR cleanup columns and quantified using a Thermo Scientific Nanodrop 8000.

For plasmids constructed by Gibson assembly, a 5 µL mixture of the protein gene and backbone was combined with 5 µL of NEB 2x Gibson assembly master mix. A DNA molar ratio of 3:1 gene:backbone was used for the Gibson assembly with a total DNA concentration of 0.1–0.5 pmol. The reaction mixture was incubated in a thermal cycler at 50°C for 1 h and then transformed into XL1-Blue *E. coli* cells (Berkeley Macrolab). After overnight growth on agar plates containing 50 µg/mL of carbenicillin for pBAD plasmids or 50 µg/mL spectinomycin for pCDF plasmids, single colonies were selected and grown overnight in 4 mL of LB medium supplemented with 50 µg/mL of the appropriate antibiotic. The plasmid was extracted from the overnight culture using a Qiagen miniprep column, and the sequence was verified using whole plasmid sequencing performed by Plasmidsaurus using Oxford Nanopore Technology with custom analysis and annotation.

For blunt end ligations, 100 ng of purified PCR mutagenesis product was combined with 2 µL 10X T4 DNA Ligase buffer, 2 µL T4 PNK (New England Biolabs), and water to 20 µL. The mixture was incubated at 37 °C for 30 min. 10 µL of the reaction was transferred to a tube containing 1 µL of T4 DNA Ligase (New England Biolabs), and ligation occurred at RT for 4-24 h. 2 µL of negative control (no ligase added) and ligation reaction were individually transformed into XL1-Blue E. coli cells (Berkeley Macrolab). Colonies were picked, grown in LB media, and sequenced as described above.

In Vivo Fluorescence Assays Using Flow Cytometry

BL21 (DE3) Star *E. coli* cells (Berkeley Macrolab) were transformed with plasmids containing S1-tagged protein, TAR RNA, or sRNA targets and were plated on LB agar plates containing respective antibiotics (carbenicillin, spectinomycin, or kanamycin each at 50 µg/mL). Plasmid combinations used in most experiments resulted in carbenicillin and spectinomycin (50 µg/mL each). At least three single colonies of each plasmid combination were used to individually inoculate 2 mL of noninducing media (NI)[6] containing respective antibiotics (each 50 µg/mL) and grown at 37°C in an incubator shaking at 250 RPM for 18-24 h until an OD600 of 3–4 was reached. The NI culture was diluted 100X into ZYP-5052 autoinduction media (AI) containing respective antibiotics (each 50 µg/mL) and grown for 18 h at 37 °C, 250 RPM in an incubator shaking to grow cells until glucose is fully consumed where lactose in the media will then express the RNA and protein.

For endpoint flow cytometry assays, 2 µL of the AI culture was diluted into 198 µL of 1X PBS buffer in a 96-well plate. Single-cell fluorescence was measured using an Attune NxT flow cytometer (Life Technologies) using the following settings: excitation lasers, 488 and 561 nm; emission channels, GFP and mCherry; cell counts for each measurement, 60,000. The data were analyzed using FlowJo software.

Orthogonal induction of TAR RNA using arabinose

BL21 (DE3) Star *E. coli* cells (Berkeley Macrolab) were transformed with pCDF-mVenus-S1 and pBAD-t-TAR (or nonbinding mutant pBAD-TAR-M1) and were plated on LB agar plates containing spectinomycin and carbenicillin (each at 50 µg/mL). At least three single colonies were individually used to inoculate 4 mL of LB media containing spectinomycin and carbenicillin (each at 50 μg/mL) in 13 mL polystyrene culture tubes and grown at 37 °C in an incubator shaking at 250 RPM for 18-24 h until an OD600 of 3–4 was reached. Fresh LB media containing spectinomycin and carbenicillin (each at 50 μg/mL) was inoculated to an OD600 of 0.1 and grown at 37°C until an OD600 of 0.3-0.6 was reached. Cultures were divided into 500 μL subcultures in 6 mL polystyrene culture tubes for induction. Subcultures were treated with 50 μM Isopropyl β-D-1-thiogalactopyranoside (IPTG), 1% arabinose, or water to account for volume changes. A low IPTG concentration (final concentration 50 μM) is required to express slower than the proteasome can degrade translated protein. The cultures continued to incubate at 37°C shaking at 250 RPM, and samples were taken over four hours. Culture samples were prepared for analysis using flow cytometry as described above.

Chemiluminescence assays for S1-tagged nanoLuciferase

BL21 (DE3) Star *E. coli* cells (Berkeley Macrolab) were transformed with pCDF-nanoLuc-S1 and pET31-t-TAR (or nonbinding mutant pET31-t-TAR-M1) and were plated on LB agar plates containing spectinomycin and carbenicillin (each at 50 μg/mL). NI and AI media cultures were prepared as described above. Using OD600 to estimate cell densities, 400-700 μL of culture was centrifuged at 4,000 *x g*, 4°C before resuspending in 500 μL 1x PBS buffer to maintain consistent total cell counts. Chemiluminescent substrate was prepared by diluting coelenterazine-h to 60 μM in reagent buffer [50 mM HEPES (pH 7.2), 100 mM KCl, 300 mM ascorbate], and equilibrating the solution at RT for at least 30 min. 96-well plates were prepared by combining 50 μL 1X PBS buffer with 50 μL resuspended cells. All nanoLuc-S1 measurements were taken at 28 °C in 1 min intervals over ten minutes in a SpectraMax i3x plate reader (Molecular Devices) after manually adding 20 μL of chemiluminescent substrate. Data shown in **Fig. S2D** was two minutes after substrate addition.
